# Stressor Identity Shapes Reactivity to Subsequent Stress

**DOI:** 10.64898/2026.09.26.754453

**Authors:** Robert J. Aukema, Lauren T. Seabrook, Byung Kwon Moon, Paloma Martinez, Claudia Klengel, Bettina Ventura, Olga Y. Ponomareva, Jakob Hartmann, William A. Carlezon, Junghyup Suh, Kerry J. Ressler

## Abstract

Extensive work has established that exposure to a single acute stressor can have profound impacts on future reactivity to new stressors, including enhanced acquisition of new fear. The impact of stressor type on development of stress sensitization, however, remains unclear. We investigated whether fundamentally different acute stressors (shock, social defeat, or restraint stress) differ in ability to sensitize acquisition of a new, contextually distinct fear memory in adult male mice two weeks following stress exposure. Stressor subtype differentially sensitized new fear learning; specifically, shock and social defeat – but not restraint stress – led to enhanced recall of a new shock-paired fear memory. We also compared how stressor subtypes differentially engage the brain-periphery interactions that regulate stress responsivity. Notably, restraint stress, which did not lead to fear sensitization to foot shock, drove similar or even greater corticosterone release than other stressors, suggesting that the neuroendocrine response to the original stressor is dissociable from development of a behavioural fear sensitization response. Using the immediate-early gene FOS as a marker of neural activity, all conditions – including the context-only control condition – activated a similar number of neurons in the basolateral amygdala. In contrast, only stressful conditions activated the paraventricular nucleus of the hypothalamus, and restraint stress selectively spared activation of the dorsal hippocampus. These data collectively suggest that future responses to stress depend critically on the identity of previously experienced stressors, rather than simply on prior exposure to stress, establishing a powerful comparative model for investigating the specific mechanisms underlying stress-enhanced fear learning.

## INTRODUCTION

A major risk for the development of post-traumatic stress disorder (PTSD) is exposure to multiple traumas experienced over time, termed ‘trauma load’ or ‘fear load’ [1–5]. Along the same lines, a major symptom of PTSD is an enhanced response to even mild stressors. Extensive work in rodents has similarly demonstrated that exposure to a single intense stressor can have profound impacts on future reactivity to new stressors, including increased avoidance behaviour in novel environments (reflective of anxiety-like behaviour) and enhanced acquisition of new fear, collectively termed sensitization [6–13]. A series of experiments, in particular from the Fanselow lab, have established the stress-enhanced fear learning (SEFL) model in rodents to investigate mechanisms linking trauma with fear sensitization [13–16]. In SEFL, exposure to repeated foot shock enhances (or “sensitizes”) fear learning to a new context paired with a single shock. This is a robust and reliable finding observed across sexes and which persists for months following the initial trauma [14,17]. Importantly, this model has strong translational validity; PTSD patients and rodent models of threat learning both display enhanced reactivity to new stressors following trauma that is resistant to exposure therapy or extinction training, respectively [13,16–19].

However, the bulk of the work investigating SEFL uses a shock stimulus both to induce stress and to assess fear learning in a new context. Thus, it is possible that fear learning is enhanced due to re-exposure to the same unconditioned stimulus (US), representing US sensitization and triggering a reminder of the original trauma. Symptoms of PTSD often occur in the absence of the original trauma, including to fundamentally different stressors; for example, interpersonal assault sensitizes response to car crash. It is thus unclear if SEFL, as originally demonstrated, is relevant to this aspect of fear sensitization or to generalization to different contexts and cue exposures. Further, there is growing evidence that certain types of trauma exposure – particularly interpersonal trauma – are associated with greater severity of PTSD symptoms [20,21], suggesting that trauma subtype may be an important moderating factor in PTSD severity.

To address these questions, we investigated whether acute stressors of distinct modality (shock, social defeat, or restraint stress) differ in ability to sensitize acquisition of a new, contextually and modality-distinct fear memory in adult male mice. We selected these specific conditions because although each of these stressors robustly activate the basolateral amygdala (BLA) [22,23], a region crucial for the development of SEFL [13], they are fundamentally different in nature – including with clear differences in sensory modality and in their conditioned responses. Understanding the underlying mechanisms linking prior trauma with subsequent fear sensitization, and how this differs by trauma subtype, has tremendous implications for the treatment, and possibly even prevention, of PTSD.

## METHODS

### Subjects

Adult male CFW mice obtained from Charles River (#024) were used as aggressors for social defeat and were pair-housed with an ovariectomized CFW female prior to experiments to maintain aggression, as previously described [24]. Adult male C57BL/6 mice obtained from Jackson Laboratory (#00064) were housed in groups of four and maintained in a temperature and humidity-controlled vivarium on a 12-hour light cycle with *ad libitum* access to food and water. All experimental procedures were approved by the Institutional Animal Care and Use Committee at McLean Hospital and follow the National Institutes of Health (NIH) Guide for the Care and Use of Laboratory Animals.

### Stress-enhanced fear learning

Animals were pre-assigned to one of four conditions (n=16 per group): no trauma (NT) control, restraint, social defeat stress (SDS), or shock. Mice were habituated once to their trauma context (Context A) and once to their new fear learning context (Context B) prior to experiments.

In phase I (**Fig. 1A**, “Trauma A”), mice were subject to stress in Context A, consisting of either restraint (60 min confined to a ventilated tube restricting all movement except breathing), SDS (3 x 5 min defeats from an aggressive resident CFW mouse over the course of an hour), shock (10 x 1 sec, 1.0 mA shocks randomly delivered over 60 min; Med Associates) [15], or a NT control condition (placed into shock chambers for 60 min without shocks delivered). All mice were returned to their group-housed home-cages following stress. 24 hours later, mice were briefly re-exposed to their respective Context A for 8 min (“Recall A)”.

**Figure 1.**
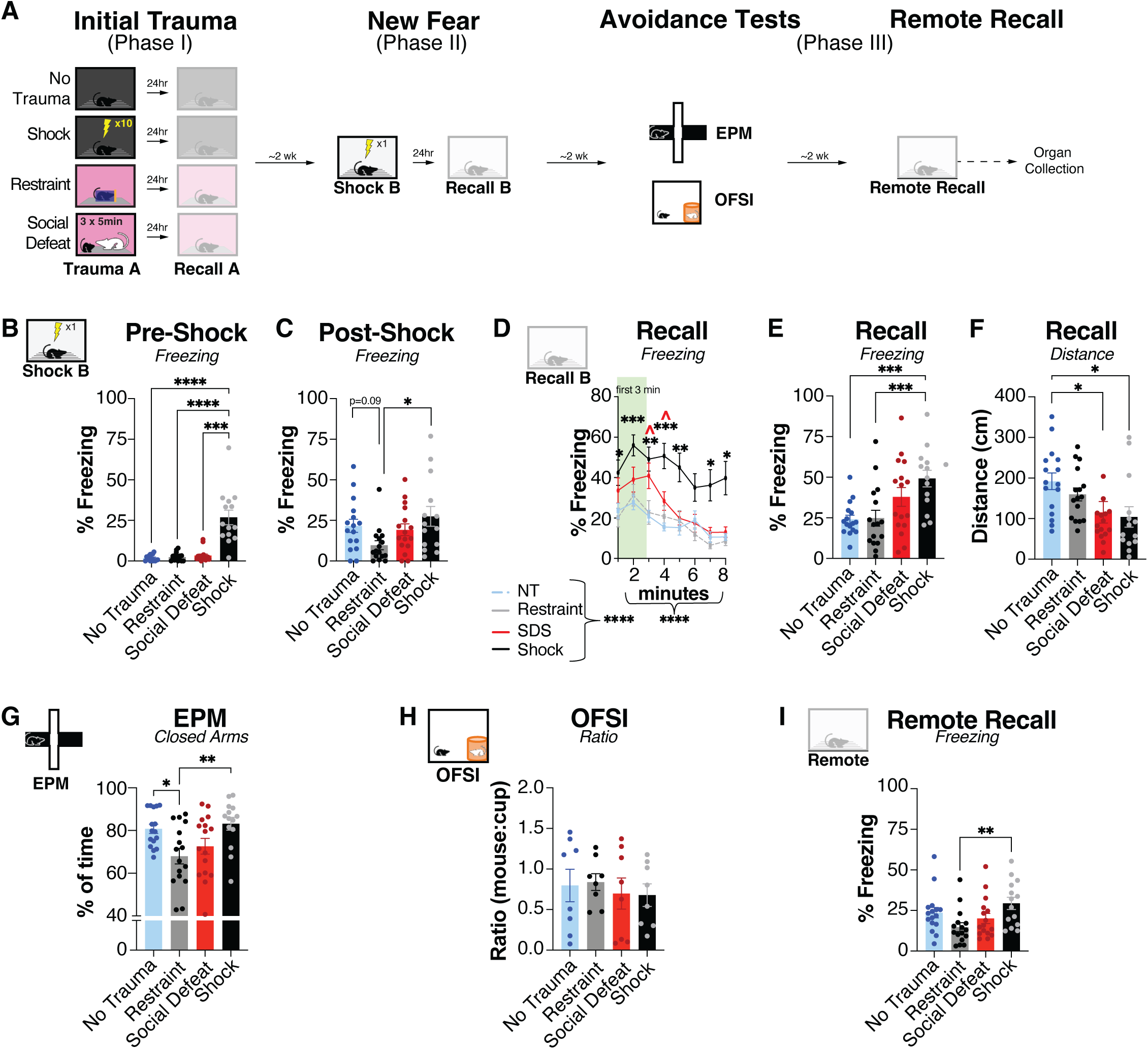
Prior trauma sensitizes new shock-paired learning in a trauma-dependent manner. **A.** Experimental timeline. **B.** Mice previously exposed to shock stress (Context A) displayed greater freezing in the baseline pre-shock period during new fear learning (Context B) than all other conditions (H_(3, 58)_ = 28.12, *p* < 0.0001; all comparisons to shock *p* < 0.001). **C.** Mice previously exposed to restraint stress displayed less freezing during the immediate 30-s post-shock period than those previously exposed to shock (H_(3, 58)_ = 9.429, *p*=0.0241; shock vs restraint: *p* = 0.0325). **D.** Mice displayed differences in freezing upon re-exposure to the new fear context depending on trauma history (Condition: F_(3, 58)_ = 11.86, *p*<0.0001; Time: F_(4.262, 247.2)_ = 18.95, *p*<0.0001; Interaction: F_(12.79, 247.2)_ = 1.306, *p* = 0.2106). **E.** History of shock increased freezing during recall of a new fear memory compared to NT controls or mice with history of restraint stress (F_(3, 58)_ = 6.129, *p* = 0.0011; NT vs shock: *p*=0.0028; restraint vs. shock: *p* = 0.0038). **F.** History of shock or SDS reduced distance traveled during recall of a new fear memory compared to NT controls (H_(3, 58)_ = 14.71, *p* = 0.0021; NT vs shock: *p*=0.0104; NT vs SDS: *p* = 0.0204). **G.** Trauma subtype altered time spent in the closed arms of the elevated plus maze (EPM; F_(3, 58)_ = 4.836, *p*=0.0045; NT vs restraint: *p* = 0.0280; restraint vs shock: *p* = 0.0088). **H.** Trauma subtype had no effect on social interaction ratio in the open field social interaction test (OFSI; F_(3, 28)_ = 0.2250, *p* = 0.8782). **I.** History of restraint stress led to reduced freezing upon remote recall of the new fear memory (H_(3, 58)_ = 11.68, *p* = 0.0086; restraint vs shock: *p*=0.0078). Data are presented as mean ± SEM. Data comparing a single measure between groups were assessed for normality using Shapiro-Wilk test and analyzed by one-way ANOVA followed by Tukey’s multiple comparisons test (for parametric data; **E, G**) or Kruskal-Wallis followed by Dunn’s test (for non-parametric data; **B, C, F, H-I**). Data comparing a single measure over time was assessed using a RM two-way ANOVA followed by Dunnett’s multiple comparisons test **(D).** Curled brace indicates main effect of condition or time. See Table S1 for full statistical results. * *p* < 0.05, ** *p* < 0.01, *** *p* < 0.001, **** *p* < 0.0001.

In phase II (**Fig. 1A**, “New Fear”), all mice were subject to a mild fear conditioning protocol in Context B two weeks following trauma [15]. This occurred in a different room than phase I, including different lighting and cleaning odorants. Mice were allowed to explore the chamber for 180 sec prior to being exposed to a single 1.0 mA, 2-sec shock delivered through the grid floor (Lafayette Instruments). After 30 sec, mice were immediately removed and returned to their home cage. The following day, mice were returned to Context B for 8 min to assess recall. Freezing was analyzed with FreezeFrame5 software (Actimetrics, Wilmette IL) and distance and location was tracked using Ethovision 14.5 software (Noldus).

In phase III (**Fig. 1A**, “Avoidance Tests”), mice were subject to testing of exploratory behaviour on the elevated plus maze (EPM) and open field social interaction (OFSI), as previously described [25]. At the termination of experiments, mice were assessed for remote recall of the new fear memory by returning to Context B for 8 min. Organs and trunk blood were collected at the termination of experiments.

### Blood, tissue, and brain collection

Adrenals and thymus were dissected from fat and weighed within 24 hours of collection. Quantification of T-cell Receptor Incision Circles (TRECs) were performed from trunk blood using qPCR as previously described, with the constant segment of C57BL/6J T-cell receptor alpha (TCRA) gene measured as reference [26]. Blood was also collected for the measurement of corticosterone (CORT). Samples were excluded if sufficient blood was not obtained within the 2 min of collection. Plasma was isolated by centrifugation at 8000 rpm for 15 min at 4 C and stored at -80 C. Corticosterone measurements were obtained using an ELISA kit (Enzo; ADI-900-097), with all samples ran in duplicate. For later time points (n=8 per group), blood was collected into EDTA-coated tubes (RAM Scientific) via tail nick at 60 min following stress onset, and via trunk blood at 90 min following stress onset. For early time points (n=8 per group), blood was collected via tail nick at 30 min following stress onset, and brains were collected following perfusions at 90 min following stress onset for quantification of FOS, as described [27]. 40 µm coronal sections of brain tissue were sectioned in a one-in-four series’ using a Leica CM-3050-S cryostat and stored in PBS with 0.02% sodium azide until processing.

### Immunohistochemistry

Immunohistochemistry for the detection of FOS was performed similar to previous experiments [28] using a rabbit anti-FOS primary antibody (cFos, Cell Signaling Technology 2250s; 1:400) and a donkey anti-rabbit secondary antibody (Alexa-488, Jackson ImmunoResearch no. 711-545-152; 1:125). Images of the BLA, PVN, or CA1 were acquired using a Zeiss Axio Imager fluorescent microscope. FOS+ neurons were automatically detected with ImageJ software, and the number of FOS+ cells were normalized to the total area and number of slices for the region of interest quantified. Animals were excluded for analyses of a given region if damage in tissue collection or processing resulted in less than two intact images (BLA: NT n=1; shock n=2; CA1: Restraint n=1; Shock n=2)

### KeyPoint Moseq Digital Behaviour Analyses

Full details are provided in Supplementary Methods. Briefly, raw .ffii videos of Context B acquired in FreezeFrame (15 fps) were exported as .avi for analyses. We labeled body parts from 600 frames using DeepLabCut (version 3.0.0) and trained using a ResNet-50-based neural network, which was used to analyze all videos in Habituation, Trauma B, Recall B, and Remote Recall days. We then applied these data to Keypoint MoSeq [29] to generate 22 behavioural “syllables”, defined as recurrent, sub-second patterns of movement. Only syllables with a frequency greater than 0.5% were included in analyses. For a broader analyses of behaviour, we combined individual syllables into 8 distinct clusters based on manually observed similarity and guided by the syllable dendrogram (**Fig. 2B**). Our output metric was syllable usage, calculated as the frequency a given syllable is employed relative to the total number of syllables used by the animal within the epoch being investigated. Analyses were restricted to only the first 3-minute epoch of Habituation and Recall B, which was then used to compare behavioural patterns across conditions. One mouse was excluded due to the model not detecting any of the top 22 syllables (shock: n=1).

**Figure 2.**
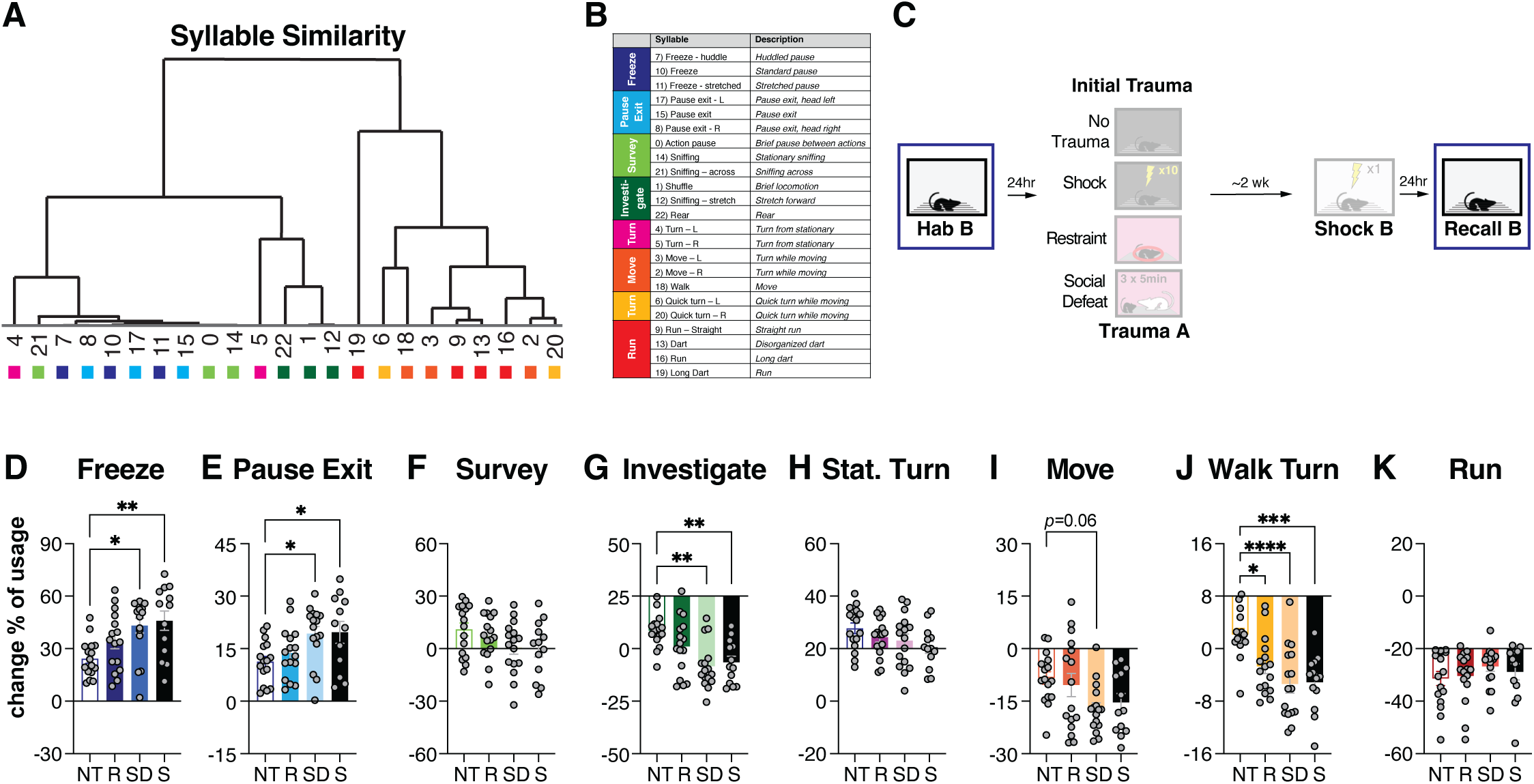
Increased freezing behaviour comes at the expense of active investigation. **A.** Syllable dendrogram, with syllables closer together in the tree displaying greater kinematic similarity. **B.** Classification of individual syllables into clusters displaying behavioural similarity. Colours of each class are displayed alongside each syllable in the syllable dendrogram to the left. **C.** Experimental timeline. Change in behaviour during the first 3 min of “HabB” compared to “Recall B” were compared between conditions. **D.** History of shock and SDS led to greater increase in freeze (H_(3, 57)_ = 13.86, *p* = 0.0031; NT vs SDS: *p*=0.0143; NT vs shock: *p*=0.0064) **E.** and pause exit usage (H_(3,57)_ = 11.73, *p*=0.0084; NT vs SDS: *p*=0.0290; NT vs shock: *p*=0.0432)**. F.** There were no differences between conditions in usage change of survey (F_(3, 57)_ = 2.036, *p* = 0.1190). **G.** History of shock and SDS led to greater decrease in usage of investigate (H_(3,57)_ = 16.65, *p*=0.0008; NT vs SDS: p=0.0015; NT vs shock: *p*=0.0072). **H.** There were no differences between conditions in usage change of stationary turn (F_(3, 57)_ = 1.828, *p* = 0.1523). **I.** History of SDS non-significantly led to greater decrease in usage of the move cluster than NT controls (F_(3,57)_ = 2.944, *p*=0.0405; NT vs SDS: *p*=0.0552). **J.** All conditions led to greater decreases in usage of walk turn compared to NT controls (F_(3,57)_ = 9.218, *p*<0.0001; NT vs restraint: *p*=0.0179; NT vs SDS: *p*<0.0001; NT vs shock: *p*=0.0004). **K.** There were no differences between conditions in usage change of run (H_(3, 57)_ = 1.041, *p* = 0.7912). Data are presented as mean ± SEM. Data were assessed for normality using Shapiro-Wilk test and analyzed by one-way ANOVA followed by Tukey’s multiple comparisons (for parametric data; **F, H-J**) or Kruskal-Wallis followed by Dunn’s test (for non-parametric data; **D-E, G, K**). See Table S1 for full statistical results. * *p* < 0.05, ** *p* < 0.01, **** *p* < 0.0001. Each data point represents an individual mouse.

### Statistics

All statistical analyses used GraphPad Prism software. Data were assessed for normality using Shapiro-Wilk test, and one-way ANOVA (followed by Tukey’s multiple comparisons) or Kruskal-Wallis test (followed by Dunn’s multiple comparisons) were performed as appropriate. Two-way ANOVA followed by Dunnett’s test to the NT condition was used to compare groups over time. A simple linear regression was performed when testing a correlation between two variables. Full statistical details are provided in **Supplementary Table 1.**

## RESULTS

### Prior trauma sensitizes new shock-paired learning in a trauma-dependent manner

We first assessed whether acute trauma sensitizes acquisition of new shock-paired fear learning – even weeks later – and whether this is modulated by trauma subtype (**Fig. 1A**). Prior to experiments, animals were habituated once to their trauma context (Context A; phase I) and once to their new fear learning context (Context B; phase II), with contexts differing by cleaning odorant, lighting, and testing room. On trauma day, animals underwent one of four trauma conditions: a “no trauma” (NT) control group that was only exposed to an empty shock chamber but no shocks were administered, shock (10 x 1.0 mA, 1 sec shocks), restraint (confined to a ventilated 50 ml Eppendorf tube with movement restricted), or social defeat stress (3 x 5 min exposures to an aggressive resident CFW mouse over 60 min). Approximately one week later, they were briefly re-exposed to their respective trauma context, without trauma exposure, to enhance reconsolidation of the trauma context [30], but otherwise remained untouched in their colony room.

Two weeks following trauma, all mice then underwent shock-paired fear learning in an entirely new context (Context B; phase II), consisting of a single, 1.0 mA, 2-sec shock. Animals previously exposed to shock trauma displayed greater basal freezing than all other groups in the 3-min period prior to shock (**Fig. 1B**). In contrast, all conditions displayed high levels of freezing immediately following shock except for animals with a history of restraint stress (**Fig. 1C**). This suggests that trauma history may differentially impact immediate reactivity to new shock stress, with history of shock increasing generalization of fear to a new shock context (i.e., increased freezing in a novel context), and history of restraint stress disrupting immediate reactivity to a new shock stress (i.e., reduced freezing immediately following new shock stress). We then returned mice to Context B, interpreting increased freezing and reduced distance traveled as reflective of greater recall. There were significant group differences in both freezing and distance traveled, particularly in the first 3 min of recall (**Fig. 1D; Supplementary Fig. 1A).** Specifically, animals with a history of shock froze more than NT controls (**Fig. 1E**). Further, history of shock or SDS also reduced distance traveled during recall of new fear (**Fig. 1F; Supplementary Fig. 1A**). Notably, animals with a history of acute restraint stress did not differ from NT controls on either measure, and in fact they froze less than those with a history of shock. Interestingly, over the entire 8 min recall, mice in the restraint condition also spent less time along the walls of the fear chambers than those with prior shock trauma (**Supplementary Fig. 1B**). Together, this suggests that trauma history may differentially alter future fear learning, with history of shock or SDS sensitizing acquisition of new shock-paired fear memory, and mice exposed to restraint stress displaying disrupted freezing behaviour in response to new shock.

We also investigated exploratory behaviour (phase III). In the elevated plus maze (EPM), mice with a history of acute restraint stress spent less time in the closed arms compared to the NT controls or those with a history of shock stress (**Fig. 1G**), suggesting increased exploration. Two days later, a subset of mice were also tested for interaction time with a novel object compared to a novel CFW mouse in the open field social interaction test (OFSI). We observed no group differences in social interaction ratio (**Fig. 1H**) and recall of new fear did not predict social interaction ratio in any group **(Supplementary Fig. 1C)**. Finally, we assessed persistence of the new fear memory by re-exposing animals to Context B four weeks after conditioning. Although there were no significant differences between groups in freezing within the first 3 min of re-exposure (data not shown), mice with a history of acute restraint stress froze less than shock-exposed animals over the entire 8 min epoch (**Fig. 1I**). Together, this suggests that history of acute restraint stress may increase exploration of novel environments and disrupt retention of newly acquired fear memories.

Chronic stress can lead to long-term changes in physiology such as increased adrenal weight and thymic involution [26]. However, we observed no differences in adrenal, thymus weight, or in plasma TRECs at the termination of experiments **(Supplementary Fig. 1D-F)**, suggesting that acute stress does not drive lasting changes in these measures as compared to prior observations with chronic stress [26].

### Increased freezing behaviour comes at the expense of active investigation

Given that history of shock or social trauma (but not acute restraint) sensitized acquisition of new shock-paired fear memory, as displayed by greater freezing and less exploration, we had two subsequent questions: (1) What other behaviours does freezing replace? and, (2) do mice with a history of restraint stress demonstrate any behavioural differences from NT controls which might not be displayed as freezing?

To examine these questions, we applied unsupervised machine learning (Keypoint-MoSeq) to videos recorded in Context B to identify broader ranges of behaviours throughout acquisition and recall of a new fear memory. This approach separated individual behaviour into recurrent, 300-400 ms patterns of movement termed ‘syllables’, which were then manually classified into broader syllable clusters based on high kinematic similarity: freeze, pause exit, passive survey, investigate, stationary turn, move, walking turn, and run (**Fig. 2A-B**). To assess how an individual animals’ baseline behaviour changes upon re-exposure to a new fear context, and whether it is influenced by trauma subtype, we next calculated the change from baseline in frequency of each syllable (**Fig. 2C**). Indeed, recall of new fear significantly altered baseline behaviour depending on syllable and trauma subtype **(Supplementary Fig. 2A-B).** Consistent with our above findings (**Fig. 1D**), we observed greater increase in usage of the “freezing” and corresponding “pause exit” syllables in mice previously exposed to shock or social defeat than NT controls (**Fig. 2D-E**). Further, we observed a parallel greater decrease in usage of the “investigate” syllable in both the SDS and shock groups (**Fig. 2G**), suggesting that increases in fear behaviour upon re-exposure to the fear context are accompanied by decreases in exploratory “active investigation”. Notably, we saw no differences between mice exposed to restraint stress and NT controls in any syllable clusters except for “walk turn”, of which a greater decrease in usage was also observed in the shock and SDS-exposed mice (**Fig. 2J**). Together, these data support our previous findings that shock and social defeat – but not restraint stress – uniquely sensitize new shock-paired fear learning, and that mice exposed to restraint stress do not display any observable behavioural differences compared to NT controls.

### Fear sensitization is not predicted by the neuroendocrine response to original trauma

Given that only some trauma subtypes (i.e., SDS and shock, but not restraint stress) led to greater expression of newly acquired fear (**Fig. 1E-F; 2D)**, we hypothesized this may be influenced by severity of the initial trauma. Social defeat stress involves natural variation in number of bites depending on the aggressor; therefore, we used total number of bites as an indirect measure of trauma severity. We observed no association in total number of bites during defeat with either freezing during recall of new fear (**Fig. 3A**) or time spent in the borders of the EPM (**Fig. 3B**). Interestingly, greater latency for defeat non-significantly predicted greater freezing during recall of new fear (**Fig. 3C**). Conversely, greater latency also non-significantly predicted less time spent in the borders of the EPM (**Fig. 3D**). This collectively suggests that rapidity of defeat may be a greater moderating factor than aggregate number of attacks, although effects are likely subtle.

**Figure 3.**
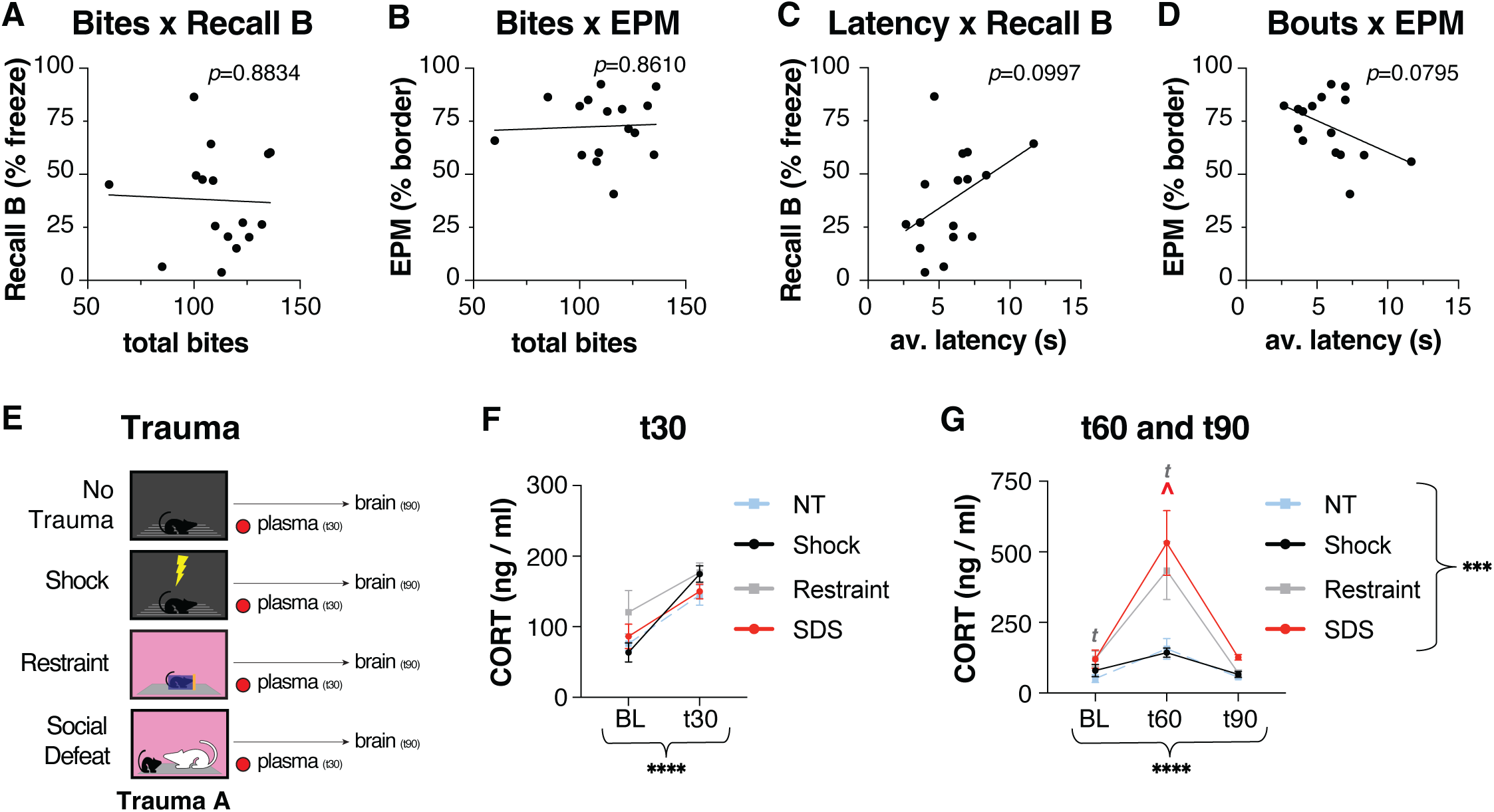
Sensitization to new stress is not predicted by HPA reactivity to the original trauma. **A.** There was no correlation between total bouts and freezing during recall of a new fear memory (r^2^ < 0.01, *p* = 0.8834) **B.** or percentage of time spent in the borders of the EPM (r^2^ < 0.01, *p* = 0.8610). **C.** There was a non-significant, positive correlation between latency of defeat and freezing during recall of a new fear memory (r^2^ = 0.18, *p* = 0.0997) **D.** and a non-significant, negative correlation between latency of defeat and percentage of time spent in the borders of the EPM (r^2^ = 0.2035, *p* = 0.0795). **E.** Experimental timeline. **F.** There was a significant effect of time in CORT measured at baseline and 30 min following trauma (Time: F_(1, 28)_ = 42.57, *p* < 0.0001; Condition: F_(3, 28)_ = 2.060, *p* = 0.1282; Interaction: F_(3, 28)_ = 1.156, *p* = 0.3438). **G.** There was a significant effect of time, condition, and an interaction between the two in CORT measured at baseline, 60-min, and 90-min following trauma (Time: F_(1.213, 29.10)_ = 37.95, *p* < 0.0001; condition: F_(3, 26)_ = 8.709, *p* = 0.0004; interaction: F_(3.368, 29.10)_ = 5.430, *p* = 0.0027; BL NT vs restraint: *p* = 0.0995; t60 NT vs restraint: *p* = 0.0970; t60 NT vs SDS: *p* = 0.0387; t90 NT vs SDS: *p* = 0.0002). Data are presented as mean ± SEM. Correlative data were tested using a simple linear regression **(A-D).** Data comparing a single measure over time were analyzed using a 2Way ANOVA or Mixed-effects analysis followed by Dunnett’s multiple comparisons test **(F-G)**. Curled brace indicates main effect of condition or time. See Table S1 for full statistical results. * *p* < 0.05, ** *p* < 0.01, *** *p* < 0.001, **** *p* < 0.0001.

We then compared the neuroendocrine response as a different measure of trauma intensity, as it is typically reflective of stressor intensity [31] and which can be directly compared across trauma subtypes. We exposed mice to 30 minutes of trauma (either shock, restraint, or social defeat stress), or exposure to a neutral no-shock context (“no trauma”) and collected plasma immediately at termination of each stressor to measure corticosterone (CORT), the main stress hormone (**Fig. 3E**). There was a main effect of time but no effect of condition or an interaction, suggesting that all stimuli similarly drove a neuroendocrine response (**Fig. 3F**). Given that CORT dynamics potentially differ between stressors, we repeated this experiment in a separate cohort of animals and exposed mice to 60 min of trauma and collected CORT 60- and 90-min following stress onset **(Supplementary Fig. 3A).** Compared to the NT control condition, restraint (*p* = 0.0970) and social defeat exposed animals had greater CORT 60 min following stress onset, which continued to be elevated in social defeat exposed animals after 90 min. Notably, there were no differences in CORT between the shock and NT condition at any timepoint. Collectively, this suggests that while each condition drove a neuroendocrine response to stress 30 min after trauma exposure, this differs in time-course between trauma subtypes and does not obviously parallel differences observed in fear sensitization. Notably, despite restraint not supporting stress sensitization in the prior experiments, it elicited the most robust neuroendocrine response among the conditions tested.

### Different trauma subtypes similarly activate the basolateral amygdala and paraventricular nucleus of the hypothalamus and differentially activate the dorsal CA1

Given a crucial role of the paraventricular nucleus of the hypothalamus (PVN) and basolateral amygdala (BLA) in stress and fear learning [31–34], and the major role of the dorsal CA1 in memory [35–37], we hypothesized that there would be differences in activation of these regions by each stressor. We measured expression of the immediate-early gene FOS protein 90 min following stress onset as an indirect marker of neural activity. Consistent with previous findings [31], all trauma subtypes increased FOS expression in the PVN compared to NT controls, with no differences between subtypes **(Supplementary Fig. 3A; Fig. 4A**). There were also no differences between trauma subtype in FOS expression in the BLA (**Fig. 4B**). Interestingly, restraint stress drove significantly less FOS expression in the CA1 than all other conditions (**Fig. 4C**). Together, this suggests that trauma subtype leads to activation patterns in a region-dependent manner, with all conditions activating the BLA, only stressful conditions activating the PVN, and restraint stress selectively sparing activation of the dorsal CA1.

**Figure 4.**
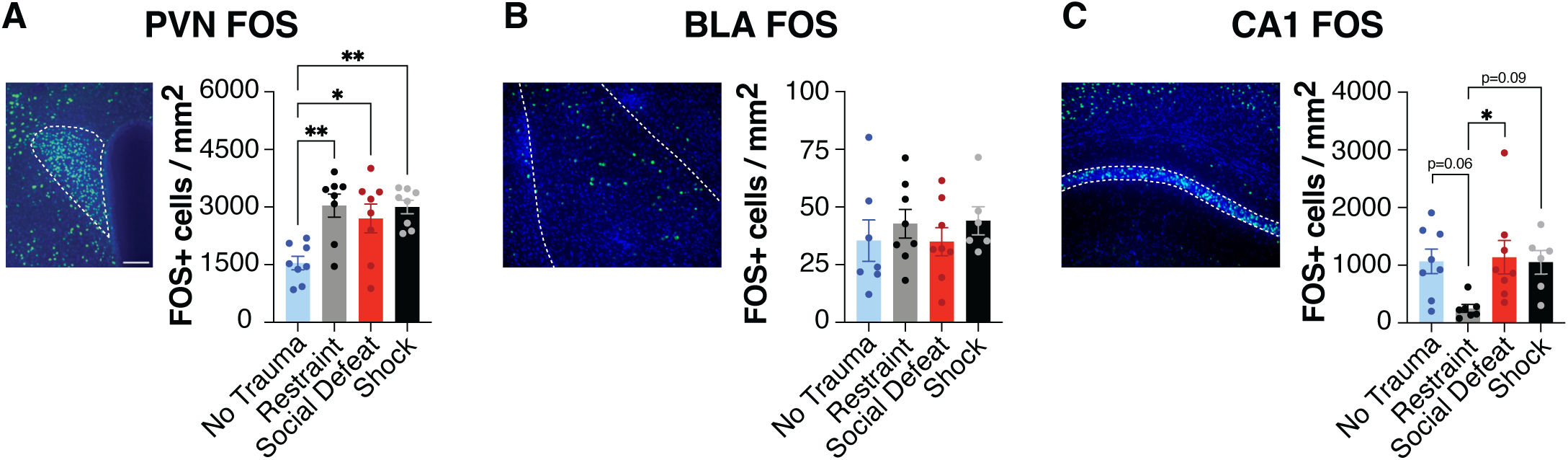
Trauma sub-types commonly activate the basolateral amygdala and differentially activate the dorsal CA1. **A.** *Left*: representative image of FOS expression in the PVN. *Right:* All stressors significantly increased FOS expression in the PVN relative to NT controls (F_(3, 28)_=6.767, *p*=0.0014; NT vs restraint: *p* = 0.0028; NT vs SDS: *p* = 0.0250; NT vs shock: *p* = 0.0035). **B.** *Left*: representative image of FOS expression in the BLA. *Right:* There were no significant differences between conditions in FOS expression in the BLA (F_(3,25)_ = 0.4629, *p*=0.7107). **C.** *Left:* representative image of FOS expression in the dorsal CA1. *Right:* Restraint stress drove significantly less FOS expression in the dorsal CA1 than all other conditions (F_(3,25)_ = 3.616, *p*=0.0270; restraint vs NT: *p* = 0.0586; restraint vs shock: *p* = 0.0348; restraint vs SDS: *p* = 0.0948). Data are presented as mean ± SEM. Data comparing a single measure were analyzed using a one-way ANOVA followed by Tukey’s multiple comparisons **(A-C).** See Table S1 for full statistical results. * *p* < 0.05, ** *p* < 0.01, *** *p* < 0.001, **** *p* < 0.0001.

## DISCUSSION

### Summary of findings

Extensive work has demonstrated that shock stress sensitizes acquisition of a new, contextually-distinct shock-paired fear memory, even weeks later [12,13,16,17]. Our work replicates these well-established findings while additionally demonstrating that a fundamentally distinct stressor from shock – namely, social defeat stress – also leads to sensitization of new shock-paired fear learning. Notably, we saw no major differences in new fear learning between mice with a history of restraint stress and NT controls, even when investigating sub-second behaviours, suggesting that acute restraint stress does not lead to lasting fear sensitization across modalities. This was surprising as restraint stress, which did not lead to fear sensitization, in fact drove a similar or even higher CORT response than other stressors. This suggests that CORT response to the initial trauma is dissociable from development of fear sensitization. Stressor subtype also led to differential activation patterns; all conditions activated the BLA, only stressful conditions activated the PVN, and restraint stress selectively spared activation of, or perhaps inhibited, the dorsal CA1. These data collectively suggest that future responses to stress depend critically on the identity of previously experienced stressors, rather than simply on prior exposure to stress, establishing a powerful comparative model for investigating the specific mechanisms underlying stress-enhanced fear learning.

### Dissociation between CORT and fear sensitization

One hypothesis for our observed behavioural differences was that stressors leading to fear sensitization may simply be more severe. We measured plasma CORT, which is typically used as an index of stressor intensity [38,39]. All stimuli increased CORT 30 min following stressor onset, including the NT control condition, consistent with previous work demonstrating that simply placing naïve animals into shock chambers induces a strong CORT response itself [39–42] (but see: [43]). At 60 min following stress, the strongest CORT responses were in the SDS and restraint conditions, consistent with restraint stress being one of the strongest laboratory stressors in driving CORT [22,44]. Yet, restraint stress did not lead to fear sensitization and further, there were no differences between NT controls and shock. This finding suggests that initial CORT response and fear sensitization are dissociable processes. Indeed, Perusini *et al* (2016) have demonstrated that although release of CORT is necessary to drive stress-enhanced fear learning, CORT itself is not sufficient for driving this phenotype [13]. Likewise, Bae *et al* (2026) have demonstrated that fear conditioning with five shocks effectively drives stress-enhanced fear learning without elevating CORT relative to NT controls [45]. In fact, blunted hypothalamic-pituitary-adrenal (HPA) axis response to stress predicts lasting stress reactivity in both rats [46] and humans [47]. As such, our findings support growing evidence that the immediate CORT response is dissociable from (or even negatively correlated with) lasting impacts of trauma.

Although typically a complementary measure to CORT response, we found PVN activation as a more reliable reflection of stress, as all stressors drove greater PVN FOS than NT controls. Given the PVN also regulates sympathetic responses to stress, likely via different neuronal populations than those regulating CORT [48,49], it is possible that heart rate may be an alternative measure of stressor intensity that could be more sensitive in capturing differences between trauma subtype. Importantly, recent work has shown that higher post-trauma heart rate and changes in heart rate variability are associated with more severe symptoms in trauma patients [50], suggesting this measure may have important predictive validity.

### Stressor subtype did not influence BLA FOS expression

We observed no differences in FOS expression within the BLA driven by any stressor subtype, even compared to the NT control condition. This was surprising, as activation of the BLA during trauma is essential for both the development of contextual fear learning [51] and stress-enhanced fear learning [13]. However, the BLA is an anatomically heterogenous structure, with high diversity among neurons in both projection target [33,52] and molecular identity [53,54], and it is also very sensitive to novelty exposure across different valences. Given that functionally distinct BLA populations support or oppose expression of fear [54–58], we therefore hypothesize that although different stressors may activate a similar total number of BLA neurons, they may activate different subpopulations [59,60]. As such, the molecular and circuit identity of stress-activated BLA neurons, and how they differ between stressor subtype, remains a crucial question.

Additionally, FOS has relatively poor temporal resolution and is thus unable to detect temporal dynamics throughout stress. Previous work using fiber photometry has demonstrated that shock and restraint stress drive different activation patterns within the BLA; restraint stress drives initially large activation of neurons which progressively decreases over time, while shock drives rapid, phasic responses, involving a progressively rising inter-shock baseline [22]. Interestingly, we observed that rapidity of defeat during SDS was a greater predictor of fear sensitization than total number of attacks, also suggesting that time-course of activation may play an important role. Divergent activation dynamics elicited by different stressors may have important consequences for synaptic strength by differentially engaging activity-dependent plasticity mechanisms. Along these same lines, previous work has shown that stressor modality can differentially influence endocannabinoid levels in the brain, including within the amygdala [61].

### Sex differences

Given known sex differences in HPA axis activation [62] and social aggression [63], our experiments focused exclusively on male mice to allow more direct comparisons between stressors. However, vast evidence has shown stress-enhanced fear learning is comparable in female mice [12,17,45,64], and we thus expect that our findings would be similarly generalizable to females although this has not been explicitly tested. Nonetheless, this is an important limitation given the higher prevalence of PTSD in women [65].

### Challenges of modeling trauma in rodents

We frequently term the first stressor (phase I) as “trauma” to distinguish from the second stressor used to drive “new fear learning” (phase II). However, modeling human trauma in rodents is challenging, as we cannot model the full complexity and context of exposures present in humans. We selected each of our trauma conditions (shock, restraint, or social defeat) because they lead to well-established and lasting impacts on behaviour or physiology in animal models [10,15], while being analogous to certain aspects of human trauma (e.g., uncontrollable; evoking a strong neuroendocrine and sympathetic response; leading to a conditioned fear response). Therefore, given that data from human populations suggests that certain subtypes of trauma may increase PTSD severity or lead to different symptom phenotypes [20,21], a major strength of our experiments lies in directly comparing different stressor or trauma subtypes.

### Conclusions & Outlook

We add to extensive human and rodent work demonstrating that prior trauma leads to lasting increases in stress reactivity [1,3,8,10–13]. Our work replicates these well-established findings while additionally demonstrating that trauma subtype is an important moderating factor in fear sensitization. Specifically, shock and social defeat stress – but not restraint stress – led to lasting fear sensitization to new shock-paired contextual fear learning. This establishes a powerful comparative model to investigate how these stressors differently impact neurobiology, potentially revealing mechanisms linking prior trauma with fear sensitization.

## CONTRIBUTIONS

Conceptualization: RJA, KJR. Experimental Design: RJA, LTS, JH, KJR. Data acquisition: RJA, LTS, PM, CK, BV, OYP, JH. Data Analysis: RJA, BKM, JS. Writing – original draft: RJA. Writing – review & editing: RJA, LTS, KJR. Funding acquisition & supervision: KJR, JS, WAC.

---

**Supplementary Figure 1.**
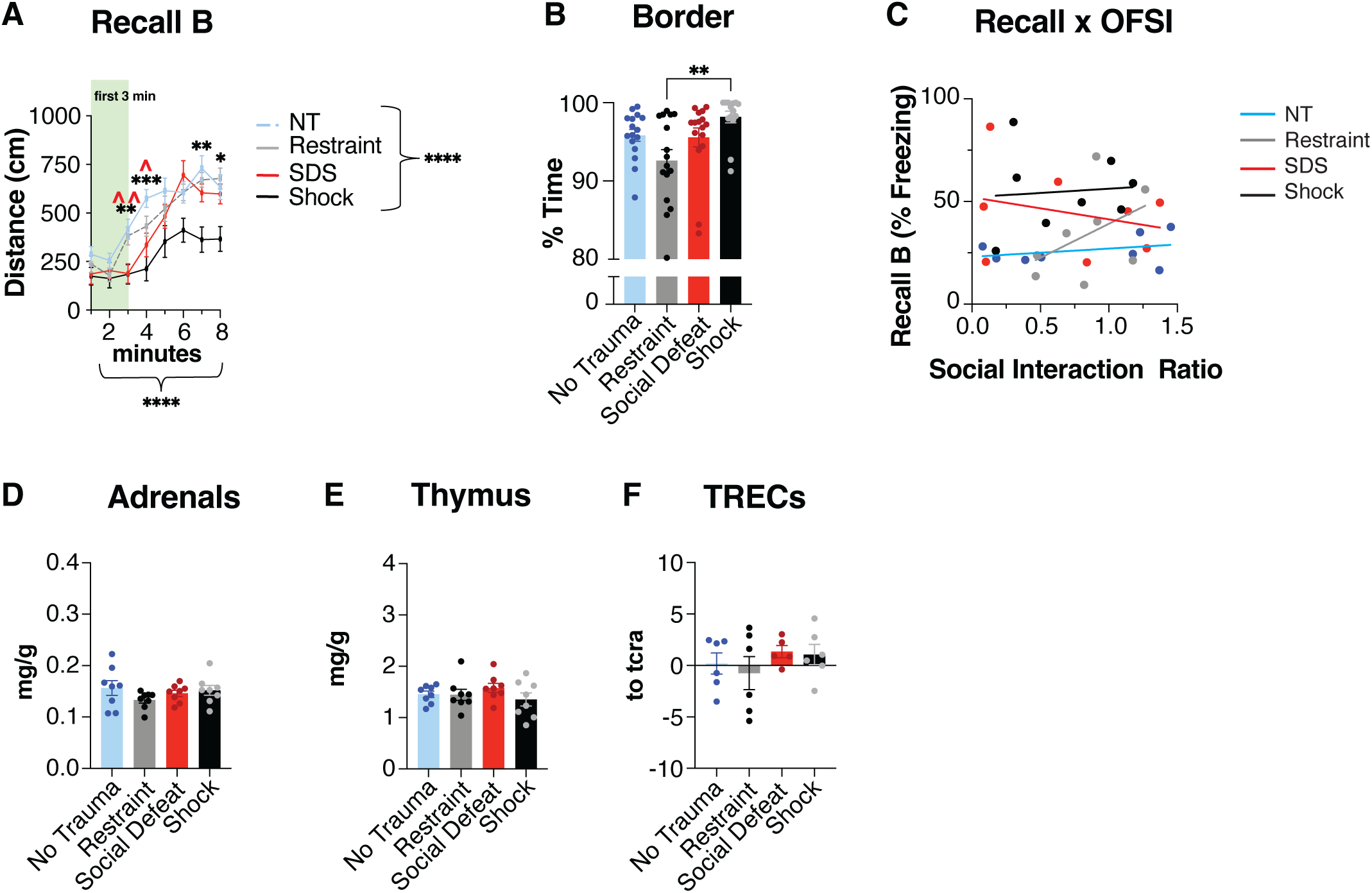
**A.** Mice displayed differences in distance traveled upon re-exposure to the new fear context depending on trauma history, time, and an interaction between the two (History: F_(3, 58)_ = 9.977, *p* < 0.0001; Time: F_(5.414, 314.0)_ = 47.25, *p* < 0.0001; Interaction: F_(16.24, 314.0)_ = 1.988, *p* = 0.0131). **B.** Mice with a history of restraint spent less time in the borders during recall B over the full 8 min compared to those with a history of shock (H_(3, 58)_ = 13.69, *p* = 0.0034; restraint vs shock: *p*=0.0017). **C.** There was no correlation between social interaction ratio in the OFSI and freezing during recall of new fear in any condition (NT: r^2^ = 0.1006, *p* = 0.4440; restraint: r^2^ = 0.2030, *p* = 0.2626; SDS: r^2^ < 0.10, *p* = 0.5310; shock: r^2^ < 0.01, *p* = 0.8427). **D.** Approximately six weeks following trauma, there were no differences in adrenal weight (F_(3, 28)_ = 1.107, *p* = 0.3628), **E.** thymus weight (F_(3, 28)_ = 0.8890, *p* = 0.4589), **F.** or plasma TRECs F_(3, 19)_ = 0.6608, *p* = 0.5863). Data are presented as mean ± SEM. Data comparing a single measure between groups were assessed for normality using Shapiro-Wilk test and analyzed by one-way ANOVA (for parametric data; **D-F**) or Kruskal-Wallis followed by Dunn’s test (for non-parametric data; **B**). Data comparing a single measure over time was assessed using a RM two-way ANOVA followed by Dunnett’s multiple comparisons test (**A**). Curled brace indicates main effect of condition or time. Correlative data was tested using a simple linear regression **(C).** See Table S1 for full statistical results. * *p* < 0.05, ** *p* < 0.01, *** *p* < 0.001, **** *p* < 0.0001.

**Supplementary Figure 2.**
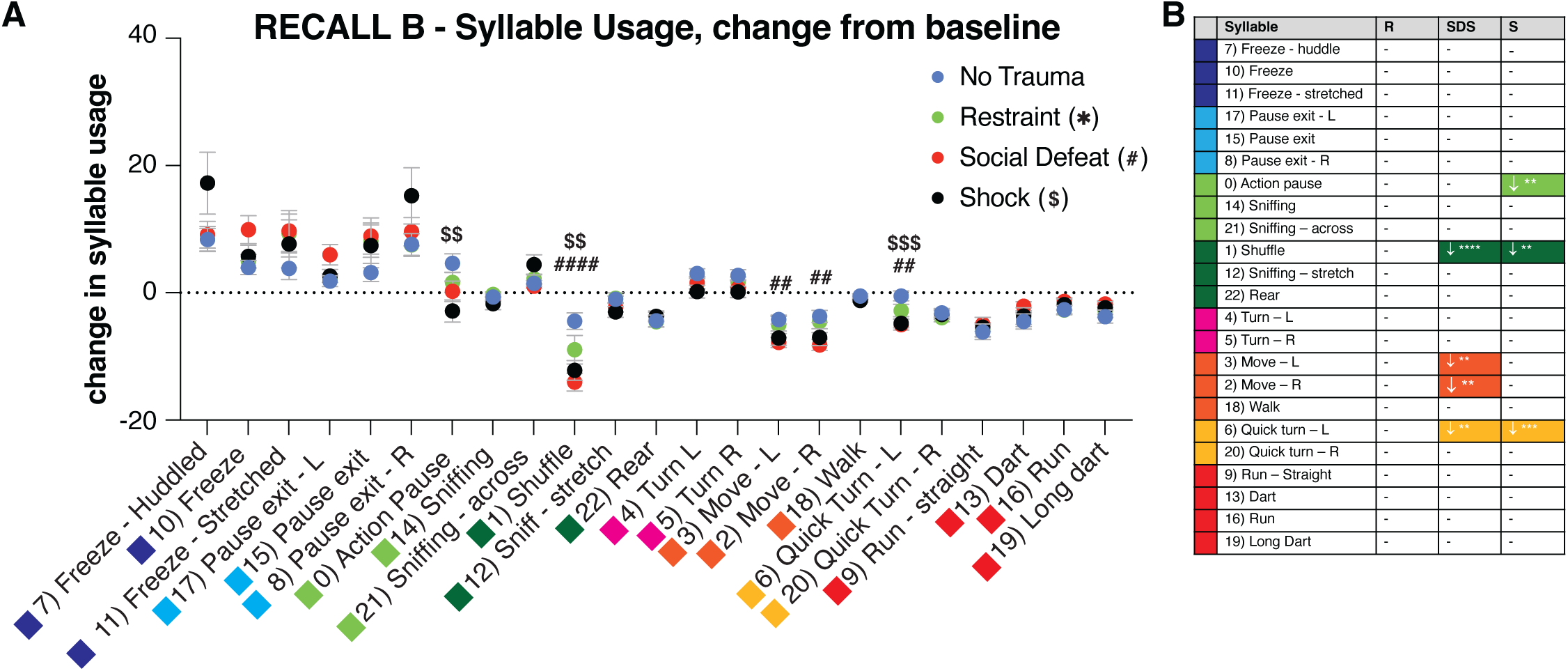
**A.** There was a significant effect of syllable and a significant interaction between syllable and condition on change from baseline in syllable usage during the first 3 min of recall of the new fear memory (Syllable: F(22, 1311) = 53.19, *p* < 0.0001; Condition: F(3, 1311) < 0.01, *p* = 0.9996; Interaction: F(66, 1311) = 2.109, *p* < 0.0001). * = significant difference between NT and restraint; ^#^ = significant difference between NT and SDS; ^$^ = significant difference between NT and shock. **B.** Summary of Dunnett’s *post-hoc* comparisons to the NT control group for each syllable. Colours correspond to broader syllable clusters identified in Figure 2A-B). Data are presented as mean ± SEM. Data comparing a single measure at multiple points were assessed using a RM two-way ANOVA followed by Dunnett’s multiple comparisons test to the NT control group (**A**). See Table S1 for full statistical results. * *p* < 0.05, ** *p* < 0.01, *** *p* < 0.001, **** *p* < 0.0001.

**Supplementary Figure 3.**
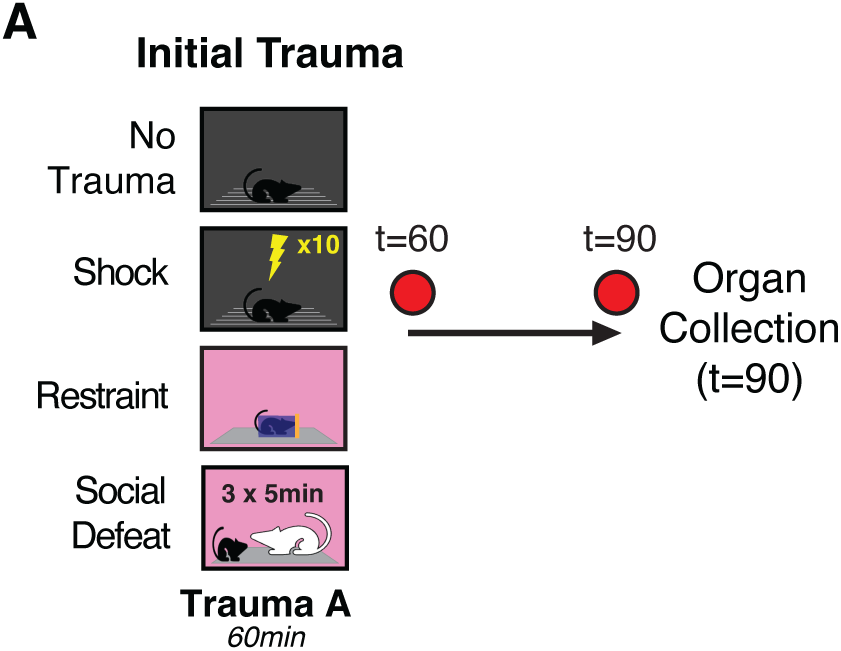
**A.** Experimental timeline.

