## Supplementary for "Stressor Identity Shapes Reactivity to Subsequent Stress"

### SUPPLEMENTARY METHODS.

#### Aggressor screening for social defeat stress.

Virgin male CFW mice were housed with ovariectomized female CFW mice for at least 1 week and screened for aggression with non-experimental C57 male mice prior to defeat sessions. Only male CFWs that showed an attack latency of less than 30 s for 2 consecutive days were used for defeats.

#### Stress-enhanced fear learning.

*Subjects.* Mice were randomly assigned to each condition however all cage-mates received the same treatment. Experiments were performed in the light phase.

*Habituation.* Animals were habituated for 20 min on separate days prior to the start of experiments. Context A differed by assigned condition. In the NT control and shock groups, this consisted of an empty fear conditioning chamber with red house lights and cleaned with 70% ethanol (Med Associates; 24 x 30 x 21 cm), with no shocks delivered. In the restraint and SDS groups, this consisted of an empty cage without access to food and water. Context B was identical for all conditions and occurred in a different behavioural room from Context A and consisted of an empty fear conditioning chamber with white house lights and cleaned with Quatricide (Lafayette Instruments; 30.5 x 25.9 x 30.5 cm).

*Trauma (Context A).* On experiment day, animals were subject to 60 min of trauma or NT control. In the NT control condition, mice were placed into a fear conditioning chamber for 60 min, lit with red house lights and cleaned with 70% ethanol (Med Associates; 24 x 30 x 21 cm), but no shock was delivered. In the shock condition, 10 x 1 sec, 1.0 mA shocks were randomly delivered through the grid floor (mean inter-shock interval = 281 sec; [15]). In the restraint condition, mice were confined to a ventilated Eppendorf tube where all movement was restricted except for breathing, and the tube containing the mouse was placed into a clean empty cage for 60 min. In the social defeat stress condition, mice were exposed to 3 x 5 min encounters with an aggressive resident CFW mouse, with a perforated PlexiGlas screen separating the mice for 15 min between each encounter and a new aggressor used each time [66]. Latency and number of bouts were recorded for each individual defeat and defeat session was terminated prior to 5 min if greater than 50 bites occurred.

*New fear learning (Context B).* Contextual fear conditioning occurred in Context B approximately two weeks following trauma, and occurred in a different room from Context A and was identical across all trauma conditions, consisting of a different conditioning chamber with a grid floor (30.5 x 25.9 x 30.5 cm, Lafayette Instruments) lit by white house lights and cleaned with Quatricide. On the day of fear learning, mice were allowed to explore the chamber for 180 sec prior to being exposed to a single 1.0 mA, 2-sec shock delivered through the grid floor. After 30 sec, mice were immediately removed and returned to their home cage. The following day (and again after 4 weeks to test remote recall), mice were returned to the same context for 8 min. Videos were recorded at 15 frames per second (fps) and freezing was analyzed with FreezeFrame5 software (Actimetrics, Wilmette IL). Thresholds to detect freezing were manually set by an experimenter blinded to conditions, with freezing defined as a lack of any observed movement for a minimum of 1 consecutive second. Videos were also exported as .avi for analyses of location and distance travelled using Ethovision 14.5 software (Noldus). One animal was excluded due to chamber malfunction during conditioning (shock: n=1) and one animal was excluded due to freezing levels during habituation exceeding 5% (shock: n=1).

**Elevated Plus Maze (EPM).**

The EPM was used to assess exploratory behaviour in a novel environment. The apparatus was elevated 70 cm above the ground and consisted of two open arms (30 x 5 cm), two closed arms (30 x 5 x 15 cm), and a central square (5 x 5 cm) with an open top. Animals were placed at the central square of the maze facing the open arm and allowed to freely explore the maze for a duration of 10 min under constant low lighting conditions. The distance traveled and zone time was tracked using Ethovision 14.5 software (Noldus).

**Open Field Social Interaction (OFSI).**

OFSI was used to measure social avoidance of a novel CFW mouse and was performed as previously described (Hisey et al, 2023). Briefly, mice were placed in a large empty arena (44 cm (w) x 44 cm (d) x 30 cm (h)) with an empty wire cup (10 cm (radius) x 20 cm (h)) for 150 s. Mice were then removed and a non-aggressive CFW was placed in the cup. Test mice were then returned to the arena for 150 s. Social interaction ratio was calculated as the amount of time spent in the 'social interaction zone' (22 cm circular zone around the wire cup) when the CFW was inside the cup divided by the amount of time spent in the zone when the cup was empty.

**Perfusions.** Brains were collected following perfusions and fixated with 4% paraformaldehyde, similar to previously described [27]: animals were anesthetized with an overdose of isoflurane and transcardially perfused with 0.9% PBS followed by 4% paraformaldehyde (PFA). Following perfusions, brains were removed and immersed in 4% PFA overnight before being switched to a 30% sucrose solution in PBS for cryoprotection. Coronal sections of 40 um were cut in four series' on a Leica CM 3050 S and stored in PBS with 0.02% sodium azide until processing.

**Keypoint Moseq.** Raw .ffii videos of Context B acquired in FreezeFrame (15 fps) were first exported as .avi for all subsequent analyses. For body part tracking we used DeepLabCut (version 3.0.0) [67–69]. We labeled 20 frames taken from 30 videos / animals for a total of 600 frames, of which 95% were used for training. We used a ResNet-50-based neural network with default parameters for 500 training iterations. We validated with a single shuffle and found the test root mean square error was 2.75, and train root mean square error was 1.98. This network was then used to analyze all videos across Habituation, Trauma B, Recall B, and Remote Recall days. Before proceeding, we manually confirmed that body parts were accurately tracked on sample videos trained in the model. We then applied these data to Keypoint MoSeq [29] and generated 22 syllables, defined as recurrent, sub-second patterns of movement. We tested out 5 different kappa values ranging $10^3$-$10^7$ to achieve a median syllable duration around 300-400 ms and proceeded with kappa=$10^5$. Using this kappa value, we ran 10 different models and chose the best model based on expected marginal likelihood (EML) score. Only syllables with a frequency greater than 0.5% were included in analyses. We then calculated the total frequency of each syllable for each trial, and a custom code was used to restrict analyses to only the first 3 minute epoch of Habituation, Recall B, and Remote Recall, and the 3 min pre-shock epoch and 32 sec post-shock epoch of Trauma B. For a broader analyses of behaviour, we combined individual syllables into 8 distinct clusters based on similarity observed manually and guided by the syllable dendrogram (**Fig. 2B**), which assigns similarity by each syllable's median trajectory. "Passive survey" was defined as stationary with head moving back and forth, and included a brief pause between actions, stationary sniffing, and sniffing left-to-right. "Active survey" was defined as stationary with body lengthening forward, and included body direction change while stationary, stretch forward, and rearing. "Freeze" was defined as a pause between surveying, and included a huddled pause, a standard pause, and a stretched pause. "Pause exit" was defined as movement from a freeze where only the head moves, and included pause exit left, pause exit, and pause exit right. "Stationary turn" was defined as a head turn only, and included survey left and survey right. "Walk" was defined as a slow, predictable locomotor activity, and included body forward right, body forward left, and quick walk. "Quick Turn" was defined as a body turn, and included walking turn right and walking turn left. "Run" was defined as a rapid change in locomotor activity, and included a straight run, a disorganized dart, a long dart, and a run. The output metric was syllable usage, calculated as the frequency a given syllable is employed relative to the total number of syllables used by the animal. This was then used to compare behavioural patterns across conditions.

### SUPPLEMENTARY TABLES

**Supplementary Table 1.** Detailed statistical test results.

| Fig | | Type of Test | Test Result | *Post hoc* test results | Notes |
|---|---|---|---|---|---|
| **MAIN FIGURES** | | | | | |
| 1 | B | Kruskal-Wallis<br>*(failed normality based on Shapiro-Wilk)* | H (3, 58) = 28.12<br>$p < 0.0001$ | Dunn's multiple comparisons<br>NT vs shock: $p < 0.0001$<br>Restraint vs shock: $p < 0.0001$<br>SDS vs shock: $p = 0.0006$ | n=14-16 per group<br><br>(n=2 excluded in shock condition due to malfunctioning chambers) |
| 1 | C | Kruskal-Wallis<br>*(failed normality based on Shapiro-Wilk)* | H (3, 58) = 9.429<br>$p = 0.0241$ | Dunn's multiple comparisons<br>NT vs restraint: $p = 0.0903$<br>Restraint vs shock: $p = 0.0325$ | n=14-16 per group<br><br>(n=2 excluded in shock condition due to malfunctioning chambers) |
| 1 | D | Two-Way RM ANOVA<br><br>Matching by time | Time:<br>F (4.262, 247.2) = 18.95<br>$p < 0.0001$<br><br>Condition:<br>F (3, 58) = 11.86<br>$p < 0.0001$<br><br>Interaction:<br>F (12.79, 247.2) = 1.306<br>$p = 0.2106$ | Dunnett's multiple comparisons to NT condition<br>min1: NT vs shock, $p = 0.0456$<br>min2: NT vs shock, $p = 0.0005$<br>min3: NT vs SDS, $p = 0.0256$<br>NT vs shock, $p = 0.0011$<br>min4: NT vs SDS, $p = 0.0476$<br>NT vs shock, $p = 0.0002$<br>min5: NT vs shock, $p = 0.0039$<br>min7: NT vs shock, $p = 0.0177$<br>min8: NT vs shock, $p = 0.0105$ | n=14-16 per group<br><br>(n=2 excluded in shock condition due to malfunctioning chambers) |
| 1 | E | Ordinary One-Way ANOVA | F (3, 58) = 6.129<br>$p = 0.0011$ | Tukey's<br>NT vs shock, $p = 0.0028$<br>Restraint vs shock, $p = 0.0038$ | n=14-16 per group<br><br>(n=2 excluded in shock condition due to malfunctioning chambers) |
| 1 | F | Kruskal Wallis<br>*(failed normality based on Shapiro-Wilk)* | H (3, 58) = 14.71<br>$p = 0.0021$ | Dunn's multiple comparisons<br>NT vs SDS: $p = 0.0204$<br>NT vs shock: $p = 0.0104$ | n=14-16 per group<br><br>(n=2 excluded in shock condition due to malfunctioning chambers) |
| 1 | G | Ordinary One-Way ANOVA | F (3, 58) = 4.836<br>$p = 0.0045$ | Tukey's<br>NT vs Restraint, $p = 0.0280$<br>Restraint vs shock, $p = 0.0088$ | n=14-16 per group<br><br>(n=2 excluded in shock condition due to malfunctioning chambers) |
| 1 | H | Ordinary One-Way ANOVA | F (3, 28) = 0.2250<br>$p = 0.8782$ | n/a | n=8 per group |
| 1 | I | Kruskal-Wallis<br>*(failed normality based on Shapiro-Wilk)* | H (3, 58) = 11.68<br>$p = 0.0086$ | Dunn's multiple comparisons<br>Restraint vs shock: $p = 0.0078$ | n=14-16 per group<br><br>(n=2 excluded in shock condition due to malfunctioning chambers) |
| 2 | D | Kruskal-Wallis | H (3, 57) = 13.86 | Dunn's multiple comparisons | |

| | | | | | |
|---|---|---|---|---|---|
| | | *(failed normality based on Shapiro-Wilk)* | $p = 0.0031$ | NT vs SDS: $p = 0.0143$<br>NT vs Shock: $p = 0.0064$ | |
| 2 | E | Kruskal-Wallis<br>*(failed normality based on Shapiro-Wilk)* | H (3, 57) = 11.73<br>$p = 0.0084$ | Dunn's multiple comparisons<br>NT vs SDS: $p = 0.0290$<br>NT vs Shock: $p = 0.0432$ | |
| 2 | F | Ordinary One-Way ANOVA | F (3, 57) = 2.036<br>$p = 0.1190$ | n/a | |
| 2 | G | Kruskal-Wallis<br>*(failed normality based on Shapiro-Wilk)* | H (3, 57) = 16.65<br>$p = 0.0008$ | Dunn's multiple comparisons<br>NT vs SDS: $p = 0.0015$<br>NT vs Shock: $p = 0.0072$ | n=13-16 per group<br><br>(n=2 excluded in shock condition due to malfunctioning chambers; n=1 due to no modeling error. |
| 2 | H | Ordinary One-Way ANOVA | F (3, 57) = 1.828<br>$p = 0.1523$ | n/a | |
| 2 | I | Ordinary One-Way ANOVA | F (3, 57) = 2.944<br>$p = 0.0405$ | Tukey's<br>NT vs SDS, $p = 0.0552$ | |
| 2 | J | Ordinary One-Way ANOVA | F (3, 57) = 9.218<br>$p < 0.0001$ | Tukey's<br>NT vs Restraint, $p = 0.0179$<br>NT vs SDS, $p < 0.0001$<br>NT vs Shock, $p = 0.0004$ | |
| 2 | K | Kruskal-Wallis<br>*(failed normality based on Shapiro-Wilk)* | H (3, 57) = 1.041<br>$p = 0.7912$ | n/a | |
| 3 | A | Simple Linear Regression | $R^2 = 0.001591$<br>$p = 0.8834$ | n/a | n = 16 |
| 3 | B | Simple Linear Regression | $R^2 = 0.002266$<br>$p = 0.8610$ | n/a | n = 16 |
| 3 | C | Simple Linear Regression | $R^2 = 0.1817$<br>$p = 0.0997$ | n/a | n = 16 |
| 3 | D | Simple Linear Regression | $R^2 = 0.2035$<br>$p = 0.0795$ | n/a | n = 16 |
| 3 | F | Two-Way RM ANOVA<br><br>Matching by time | Time: F (1, 28) = 42.75<br>$p < 0.0001$<br><br>Condition:<br>F (3, 28) = 2.060<br>$p = 0.1282$<br><br>Interaction:<br>F (3, 28) = 1.156<br>$p = 0.3438$ | Dunnett's multiple comparisons to NT condition | n = 8 per group |
| 3 | G | Mixed Effects Analysis<br><br>Matching by time | Time:<br>F (1.213, 29.10) = 37.95<br>$p < 0.0001$<br><br>Condition:<br>F (3, 26) = 8.709<br>$p = 0.0004$ | Dunnett's multiple comparisons to NT condition<br><br>BL: NT vs Restraint, $p = 0.0995$<br>t60: NT vs Restraint, $p = 0.0970$<br>NT vs SDS, $p = 0.0387$<br>t90: NT vs SDS, $p = 0.0002$ | n = 6-8 per group<br><br>(n=2 animals missing in NT condition due to too long of collections at t60, thus causing stress; n=3 samples in restraint and n=1 samples in SDS |

| | | | | | |
|---|---|---|---|---|---|
| | | Geisser-Greenhouse correction | Interaction:<br>F (3.638, 29.10) = 5.430<br>*p* = 0.0027 | | missing due to not enough plasma collected) |
| 4 | A | Ordinary One-Way ANOVA | F (3, 28) = 6.767<br>*p* = 0.0014 | Tukey's<br>NT vs restraint: *p* = 0.0028<br>NT vs SDS: *p* = 0.0250<br>NT vs shock: *p* = 0.0035 | n=8 per group |
| 4 | B | Ordinary One-Way ANOVA | F (3, 25) = 0.4629<br>*p* = 0.7107 | n/a | n=6-8 per group<br>(n=1 missing in NT and n=2 missing in shock due to damaged tissue) |
| 4 | C | Ordinary One-Way ANOVA | F (3, 25) = 3.616<br>*p* = 0.0270 | Tukey's<br>NT vs restraint: *p* = 0.0586<br>Restraint vs SDS: *p* = 0.0348<br>Restraint vs shock: *p* = 0.0948 | n=6-8 per group<br>(n=1 missing in restraint and n=2 missing in shock due to damaged tissue) |
| **SUPPLEMENTARY FIGURES** | | | | | |
| 1 | A | Two-Way RM ANOVA<br><br>Matching by Time | Time:<br>F (5.414, 314.0) = 47.25<br>*p* < 0.0001<br><br>Condition:<br>F (3, 58) = 9.977<br>*p* < 0.0001<br><br>Interaction:<br>F (16.24, 314.0) = 1.988<br>*p* = 0.0131 | Dunnett's multiple comparisons to NT condition<br>min3: NT vs SDS, *p* = 0.0098<br>NT vs shock, *p* = 0.0092<br>min4: NT vs SDS, *p* = 0.0114<br>NT vs shock, *p* = 0.0002<br>min5: NT vs shock, *p* = 0.0505<br>min6: NT vs shock, *p* = 0.0680<br>min7: NT vs shock, *p* = 0.0012<br>min8: NT vs shock, *p* = 0.0161 | n=14-16 per group<br><br>(n=2 excluded in shock condition due to malfunctioning chambers) |
| 1 | B | Kruskal-Wallis *(failed normality based on Shapiro-Wilk)* | H (3, 58) = 13.69<br>*p* = 0.0034 | Dunn's multiple comparisons<br>NT vs shock, *p* = 0.0703<br>Restraint vs Shock, *p* = 0.0017 | |
| 1 | C | Simple Linear Regression | NT:<br>$R^2$ = 0.1006<br>*p* = 0.4440<br><br>Restraint:<br>$R^2$ = 0.2030<br>*p* = 0.2626<br><br>SDS:<br>$R^2$ = 0.06856<br>*p* = 0.5310<br><br>Shock:<br>$R^2$ = 0.007106<br>*p* = 0.8427 | n/a | n=8<br><br><br><br>n=8<br><br><br><br>n=8<br><br><br><br>n=8 |
| 1 | D | Ordinary One-Way ANOVA | F (3, 28) = 1.107<br>*p* = 0.3628 | n/a | n=8 per group |
| 1 | E | Ordinary One-Way ANOVA | F (3, 28) = 0.8890<br>*p* = 0.4589 | n/a | n=8 per group |

| | | | | | |
|---|---|---|---|---|---|
| 1 | F | Ordinary One-Way ANOVA | F (3, 19) = 0.6608<br>$p = 0.5863$ | n/a | n=5-6 per group<br><br>(n=2 in NT, n=2 in restraint, n=3 in SDS, n=2 in shock missing due to insufficient blood) |
| 2 | A | Two-Way RM ANOVA<br><br>Matching by syllable | Syllable:<br>F (3.215, 183.2) = 50.88<br>$p < 0.0001$<br><br>Condition:<br>F (3, 57) = 3.496<br>$p < 0.0001$<br><br>Interaction:<br>F (9.644, 183.2) = 2.017<br>$p = 0.2106$ | Dunnett's multiple comparisons to NT condition<br>Syl00: NT vs shock, $p = 0.0087$<br>Syl01: NT vs SDS, $p < 0.0001$<br>NT vs shock, $p = 0.0024$<br>Syl02: NT vs SDS, $p = 0.0045$<br>Syl03: NT vs SDS, $p = 0.0055$<br>NT vs shock, $p = 0.0969$<br>Syl04: NT vs shock, $p = 0.0872$<br>Syl06: NT vs SDS, $p = 0.0014$<br>NT vs shock, $p = 0.0009$<br>Syl10: NT vs SDS, $p = 0.0625$<br>Syl17: NT vs SDS, $p = 0.0809$ | n=13-16 per group<br><br>(n=2 excluded in shock condition due to malfunctioning chambers; n=1 due to no modeling error. |

**Note:** *post-hoc* values only reported if $p < 0.10$
